# An Open-Source Membrane Stretcher for Simultaneous Mechanical and Structural Characterizations of Soft Materials and Biological Tissues

**DOI:** 10.1101/2024.05.01.592106

**Authors:** Shannon Li, Alyssa Gee, Nathan Cai, Alexandra Bermudez, Neil Y.C. Lin

## Abstract

The ability to simultaneously measure material mechanics and structure is central for understanding their nonlinear relationship that underlies the mechanical properties of materials, such as hysteresis, strain-stiffening and -softening, and plasticity. This experimental capability is also critical in biomechanics and mechanobiology research, as it enables direct characterizations of the intricate interplay between cellular responses and tissue mechanics. Stretching devices developed over the past few decades, however, do not often allow simultaneous measurements of the structural and mechanical responses of the sample. In this work, we introduce an open-source stretching system that can apply uniaxial strain at a submicron resolution, report the tensile force response of the sample, and be mounted on an inverted microscope for real-time imaging. Our system consists of a pair of stepper-based linear motors that stretch the sample symmetrically, a force transducer that records the sample tensile force, and an optically clear sample holder that allows for high-magnification microscopy. Using polymer samples and cellular specimens, we characterized the motion control accuracy, force measurement robustness, and microscopy compatibility of our stretching system. We envision that this uniaxial stretching system will be a valuable tool for characterizing soft and living materials.

## Specifications table

|  |  |
| --- | --- |
| **Hardware name** | Stretchoscope – a membrane stretcher for simultaneous mechanical and structural characterizations of soft materials and biological tissues |
| **Subject area** | Bioengineering, Mechanical Engineering, and Material Science |
| **Hardware type** | Mechanical Testing and Imaging |
| **Closest commercial analog** | *Flexcell* Cell Stretching Bioreactors, Cultureware, Curi Bio Cytostretcher |
| **Open source license** | CC-BY 4.0 |
| **Cost of hardware** | ~ USD \$4,000 |
| **Source file repository** | Within the article |

## 1. Hardware in context

Characterizing the mechanical properties and structural rearrangements of both living materials (e.g., cells and tissues), and soft materials (e.g., polymers), is crucial for understanding each system’s response to mechanical stimuli, which essentially govern cell behavior or the material’s physical properties, respectively. Specifically, it has been demonstrated that mechanical cues play a pivotal role in various cellular processes [12, 7], including migration, differentiation, and tissue development while mechanical forces exerted on soft, non-living materials can alter chain alignment, intermolecular interactions, and introduce material defects and stress concentrations [15, 9].

Over the last few decades, many devices have been developed to conduct integrated structural and mechanical measurements for soft materials. For example, parallel-plate shear cells[17, 25, 6, 5] can characterize the structural and mechanical responses of sheared complex liquids. However, these devices are typically limited to the characterizations of the shear response of viscoelastic fluids. For cellular materials, most devices focus on the capability of stretching and imaging mechanically perturbed cells [14, 13, 11], omitting the force measurement. There has also been development of open-source stretchers [13, 19, 18, 20, 24] to make the cell stretching systems more accessible. Recently, there have been developments of cell stretchers that also allow for force measurements [22]. For example, the deformation of a silicone sheet has been used to approximate the elastic module of a cell layer [22]. The bending of rods has been used to infer the force imposed on partially digested cell monolayers [10]. The impact of fiber alignment and depth in articular cartilage has been inspected through shear testing and elastography [21, 2]. Uniaxial characterization has been done on ultra-thin freestanding films [3]. Low cost devices have also been designed for applying uniaxial strain to cells [1]. Despite these instrumental advancements, the field still lacks a device that can characterize the structural rearrangements of various membrane samples while directly quantifying the system’s mechanical response.

To address this technological gap, we engineered a uniaxial stretching device – stretchoscope [4]. Our stretchoscope is integrated with microscopy and a force transducer to enable live imaging and acquisition of stress response data, compatible with both living and physical materials. This integrative open-source platform allows for precise control of mechanical deformation while simultaneously providing high-resolution imaging of the material’s microstructure for real-time characterization and quantitative measurements of mechanical properties, offering a comprehensive approach that bridges the gap between structural characterization and mechanical testing. We envision our stretchoscope can be a versatile tool to advance the fields of mechanobiology and material science by providing a platform enabling biologists to unravel the complex mechanistic principles underlying mechanotransduction-driven cellular behaviors, and by enabling material scientist to investigate structure-property relationships, deformation mechanisms, and failure modes of these materials for downstream applications in fields such as biomedical instrumentation, flexible electronics, and soft robotics.

## 2. Hardware Description

The stretchoscope hardware comprises two stepper motors that are attached to a stainless steel base plate and move uniaxially in opposing directions to stretch a sample mounted to the motor arms (Fig. 1a). The motor arms suspend the sample over a transparent imaging window in the center of the base plate to enable simultaneous stretching and sample imaging (Fig. 1b). The current design builds upon past prototypes of similar components by utilizing closed-loop stepper motors that can apply force, in which the inner diameter of the imaging window allows a maximum of *∼* 80% tensile strain to be applied to the sample. We also demonstrate this model can be accessibly constructed by three-dimensional (3D) printing the mechanical parts (Fig. 1c), in which all of the design files have been made publicly available [16]. In the following sections, we introduce the three major parts of the stretchoscope hardware: (1) motor arms that connect the sample, force transducer, and stepper motors, (2) the imaging window that allows for inverted high-magnification microscopy, and (3) jig components that secure the membrane samples near the imaging window. We also discuss how these parts can be manufactured using hobbyist grade fused deposition modeling (FDM) 3D printers.

**Figure 1:**
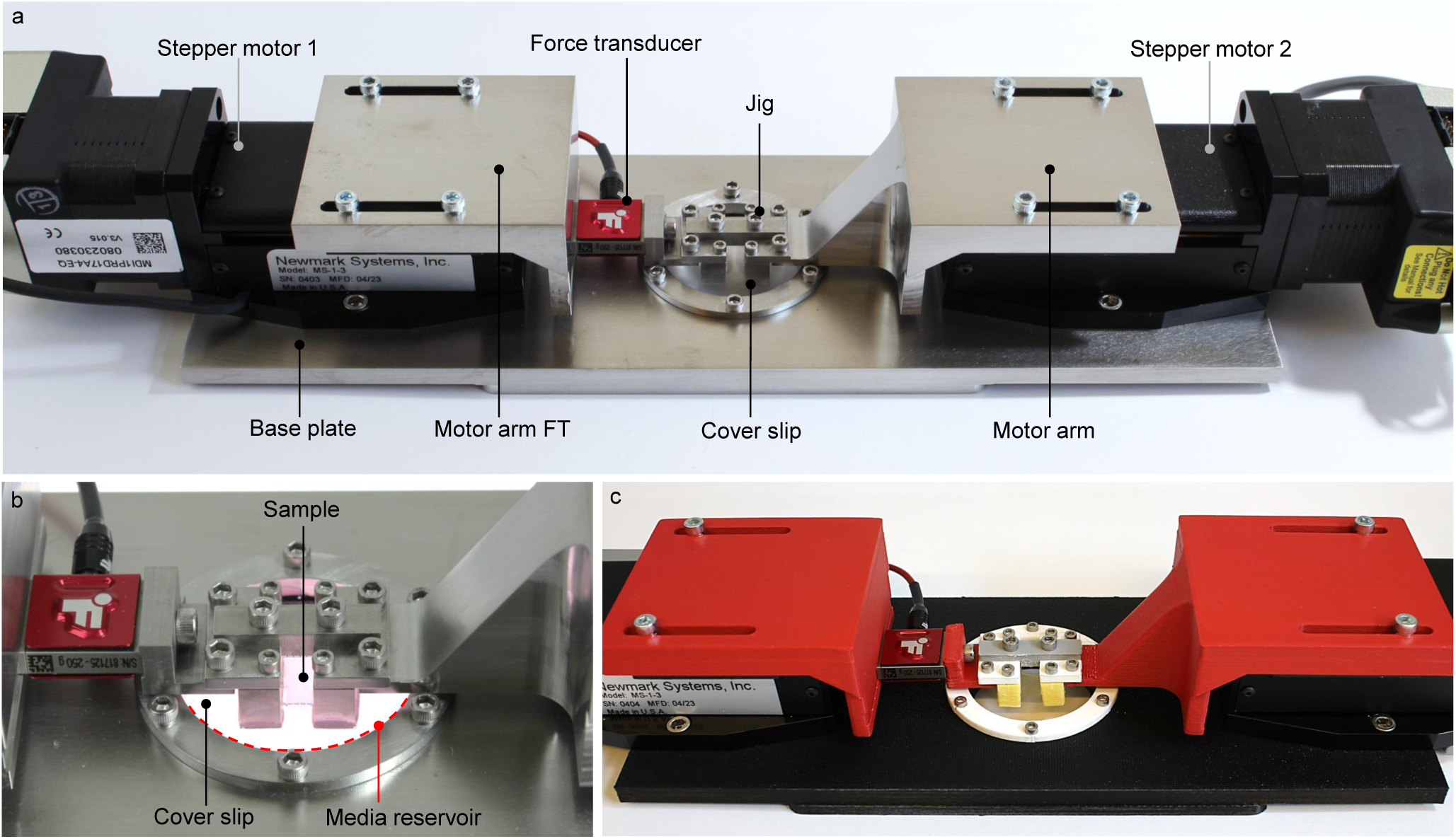
Photos overviewing the design and assembly of the stretchoscope. (a) Overview of a complete assembly of the stretchoscope that is comprised of machined mechanical parts. (b) Enlarged view of the imaging window with a transparent PDMS polymer sample submerged in media. The media is in a reservoir supported by a glass cover slip in the viewing window. (c) A complete assembly of stretchoscope that is comprised of 3D-printed mechanical parts.

### 2.1 Motor arms

The motor arms are designed to serially connect the sample, force transducer, and stepper motors, enabling sample stretching and force measurements. The stretchoscope motor arms have two types. One type of motor arm directly connects the motor to the jig, where the sample is mechanically secured to it using screws. The other type of motor arm is the force transducer (FT) motor arm. This arm is composed of two parts in which one part connects the motor to the force transducer and the other joins the force transducer to the jig (Fig. 1a). If the experiment does not require the force measurement output, the single part motor arm first described can be installed in place of the motor arm FT.

### 2.2 Imaging window and media reservoir

The circular cutout in the base plate is multi-functional in that it allows the sample to be imaged while also acting as a media reservoir for biological samples. This imaging window contains a circular glass coverslip, a silicone O-ring, and a metal cover ring which mechanically compresses the layers together (Fig. 2a). Together, these components form a leak-proof liquid chamber to allow the samples to be fully immersed in the appropriate medium during experiments. For example, this system preserves the water content of hydrogel samples as well as the physiological condition (e.g., cell culture media) of living biological specimens. The imaging window dimensions are optimized to accommodate small samples while allowing high numerical aperture (NA) microscope objectives, which typically have a short working distance, to reach the coverslip. For long term imaging of biological samples spanning several hours or days, the appropriate culture conditions can be maintained by directly mounting the stretcher onto a microscope housed in an incubator, such as an Incucyte or Etaluma microscope. It is recommended to increase the humidity of the incubator for multi-day acquisition to minimize media evaporation. Alternatively, users may also house the stretcher in an onstage incubator if an in-incubator microscope is inaccessible.

**Figure 2:**
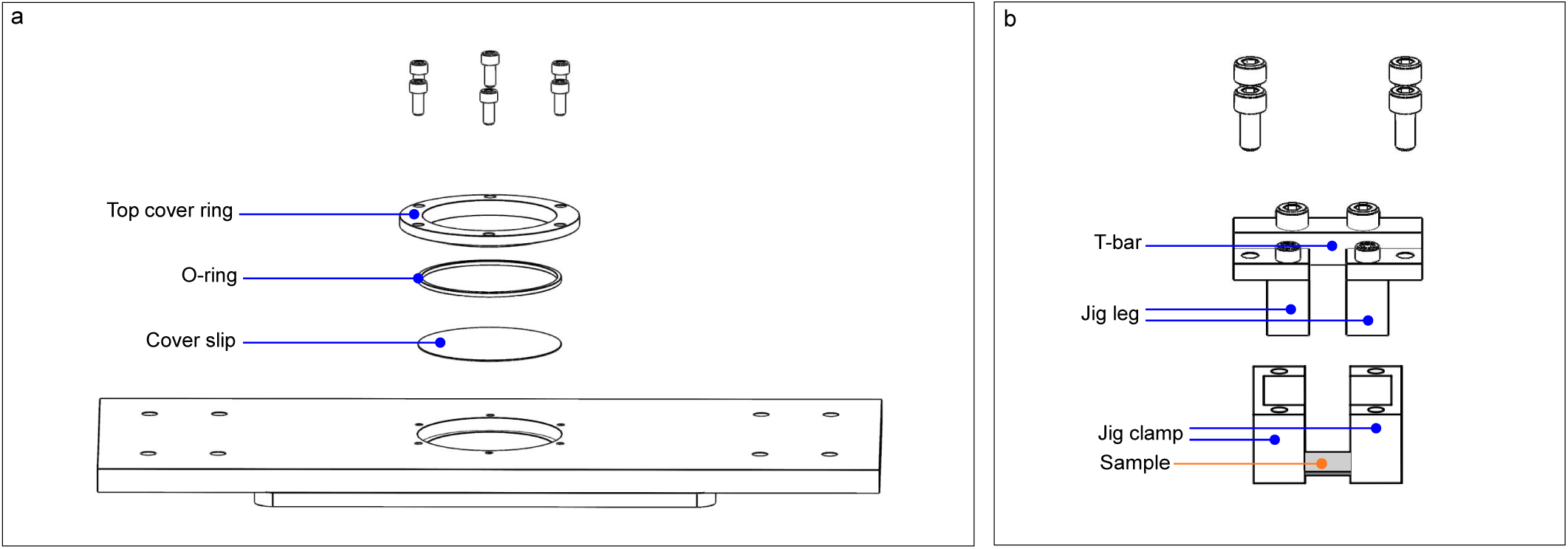
(a) Exploded view of the layers comprising the imaging window. (b) Exploded view of the jig assembly. The location of the mounted sample is annotated in orange while assembly components are annotated in blue. For clarity, the exploded view of the T-bar and jig legs is shown in Fig. 4b.

### 2.3 Jig

The jig component is designed to provide a low-profile mount of the sample near the imaging window. To maintain sample integrity and sample alignment during mounting, we utilize a T-bar that connects the two jig legs such that they remain a fixed distance apart from one another. Using a T-bar during sample fabrication and mounting reduces sample slacking since the fabricated samples always maintain their initial dimensions. Before stretching, the T-bar must be removed to allow the jig legs to be pulled apart by the motors. For mounting tissues or inert samples, the fixed jig legs have corresponding jig clamps that slide up the legs to securely hold the sample in place and avoid sample slipping during stretching (Fig. 2b). Sample tautness can also be manually tuned by first securing one end of the sample and then gently stretching the unclamped sample-end to the desired tautness before securing with the other clamp. By holding the membrane samples to the imaging window within *∼* 1 mm, high-magnification and high-NA microscope objectives, such as Nikon CFI Fluor 60x water immersion objective, can be used to image the microstructure of the samples, providing a *∼* 250 nm lateral resolution when conducting fluorescent microscopy.

### 2.4 3D printed parts

We demonstrate that the manufactured metal parts can instead be 3D-printed as a more accessible and cost-effective option (Fig. 1c). Using Prusa i3 MK3S+ 3D printers, we printed all the components forming the motor arms, imaging window, and jig system with 30% infill and 0.2 mm layer height. Settings may be adjusted for better printing resolution and tolerance. The screw hole dimensions are decreased by 10% so that the screws can be threaded in accordingly.

## 3. Design files summary

*Repository for Files: Part Files*

| Design File Name | Description | File type | Open source license | File location |
| --- | --- | --- | --- | --- |
| BasePlate | Base plate of stretcher | STEP | CC-BY 4.0 | URL |
| MotorArm | Non-force transducer motor arm | STEP | CC-BY 4.0 | URL |
| MotorArmFT_Pt1 | Motor arm between motor & force transducer | STEP | CC-BY 4.0 | URL |
| MotorArmFT_Pt2 | Motor arm between jig & force transducer | STEP | CC-BY 4.0 | URL |
| TopCoverRing | Top cover ring keeps coverslip in place | STEP | CC-BY 4.0 | URL |
| JigLegs | Side of jig for sample suspension | STEP | CC-BY 4.0 | URL |
| JigClamp | Jig clamp for securing sample to jig legs | STEP | CC-BY 4.0 | URL |
| JigTBar | Jig T-bar for holding jig together | STEP | CC-BY 4.0 | URL |
| Control_Interface_Main | Control interface LabVIEW file | VI | CC-BY 4.0 | URL |

## 4. Bill of materials summary

| Designator | Component | Qty | Unit cost (USD) | Total cost (USD) | Material source | Part Number | Material type |
| --- | --- | --- | --- | --- | --- | --- | --- |
| Motor | Stepper motor | 2 | \$1,447.00 | \$2,894.00 | Newmark Systems | MS-1-3 | Electronics |
| Force transducer | Force transducer | 1 | \$848.00 | \$848.00 | FUTEK | FSH04401 | Electronics |
| Force transducer connector | USB 220 connector | 1 | \$1,100.00 | \$1,100.00 | FUTEK | FSH04720 | Electronics |
| Motor cable | MDrive RS422 ribbon cables | 2 | \$70.00 | \$140.00 | Newmark Systems | 250168 | Electronics |
| Cable | Serial to USB converter | 2 | \$25.88 | \$51.76 | Gearmo | US-485422 | Electronics |
| Power adapter | Power adapter | 2 | \$17.88 | \$35.76 | Gonine | JSJ-AC211197G | Electronics |
| Power hub | USB hub | 1 | \$8.95 | \$8.95 | SABRENT | HB-UMLS | Electronics |
| O-ring | 2.65 x 35.5 mm rubber O-ring | 1 | \$0.41 | \$0.41 | McMaster-Carr | 3799N38 | Rubber |
| Coverslip | No. 1.5 glass coverslips, 40 mm round | 1 | \$0.20 | \$0.20 | Warner Instruments | CS-40R15 | Glass |
| Fasteners | 4-40 x 1/4" socket head cap screws | 10 | \$0.06 | \$0.60 | McMaster-Carr | 92196A106 | Stainless Steel |
| Fasteners | 8-32 x 3/16" socket head cap screws | 4 | \$0.06 | \$0.24 | McMaster-Carr | 92196A189 | Stainless Steel |
| Fasteners | 6-32 x 5/16" socket head cap screws | 8 | \$0.07 | \$0.56 | McMaster-Carr | 92196A145 | Stainless Steel |
| Fasteners | 2-56 x 1/4" socket head cap screws | 4 | \$0.08 | \$0.32 | McMaster-Carr | 92196A077 | Stainless Steel |

## 5. Build instructions

### 5.1 Motors and imaging window

Parts required:

(*×*1) Base plate

(*×*2) Stepper motor

(*×*1) Coverslip

(*×*1) Silicone O-ring (2.65 mm thick, 35.5 mm diameter)

(*×*1) Top cover ring

(*×*4) 8-32 x 3/15” socket head cap screw

(*×*6) 4-40 x 1/4” socket head cap screw

Screw the stepper motors to the base plate using 8-32 socket head cap screws. The screws will go through the side panels on the bottom sides of the motors and into the pair of screw holes closest to the short edge of the base plate on opposite sides (Fig. 3, red arrows). For the circular viewing window, first carefully place a clean glass cover-slip into the hole on the base plate, then fit the O-ring into the hole, making sure the edges are flush with the metal sides and in contact the cover-slip below. Place the metal cover ring on top. Line up the screw holes on the ring to the base plate and secure the layers using 4-40 screws (Fig. 3, blue arrows). No additional components are required to secure the coverslip. The screws into the metal ring cover provide enough compression to hold all components of the imaging window securely in place.

**Figure 3:**
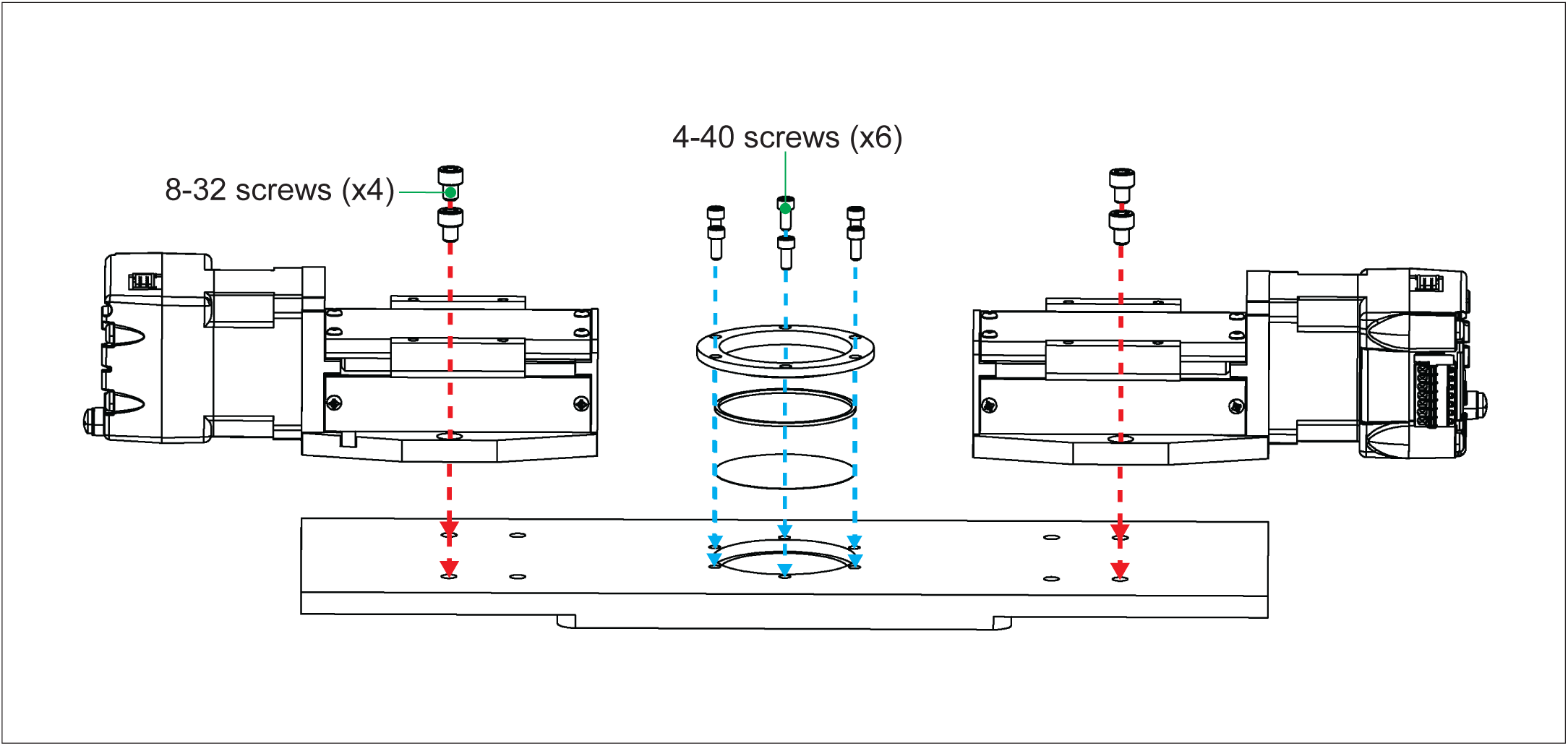
Steps to attach stepper motors and imaging window parts to base plate.

### 5.2 Force transducer and jig assembly

Parts required:

(*×*1) Motor arm part 1

(*×*1) Motor arm part 2

(*×*1) Force transducer

(*×*2) Jig leg

(*×*2) Jig clamp

(*×*1) Jig T-bar

(*×*2) M3 socket head cap screw

(*×*4) 2-56 socket head cap screw

Separate from the base assembly, attach the force transducer in between the force transducer motor arm part 1 and part 2 using M3 screws (Fig. 4a, red arrows). The silver part of the force transducer should sit into the slots of the motor arm parts. For the jig, connect the T-bar to the jig legs using 4-40 screws (Fig. 4b, blue arrows). Use 2-56 screws to slot the jig clamps into the jig legs with the open edge facing inwards (Fig. 4b, red arrows). Mounting a sample to the jig with jig clamps will be discussed later.

**Figure 4:**
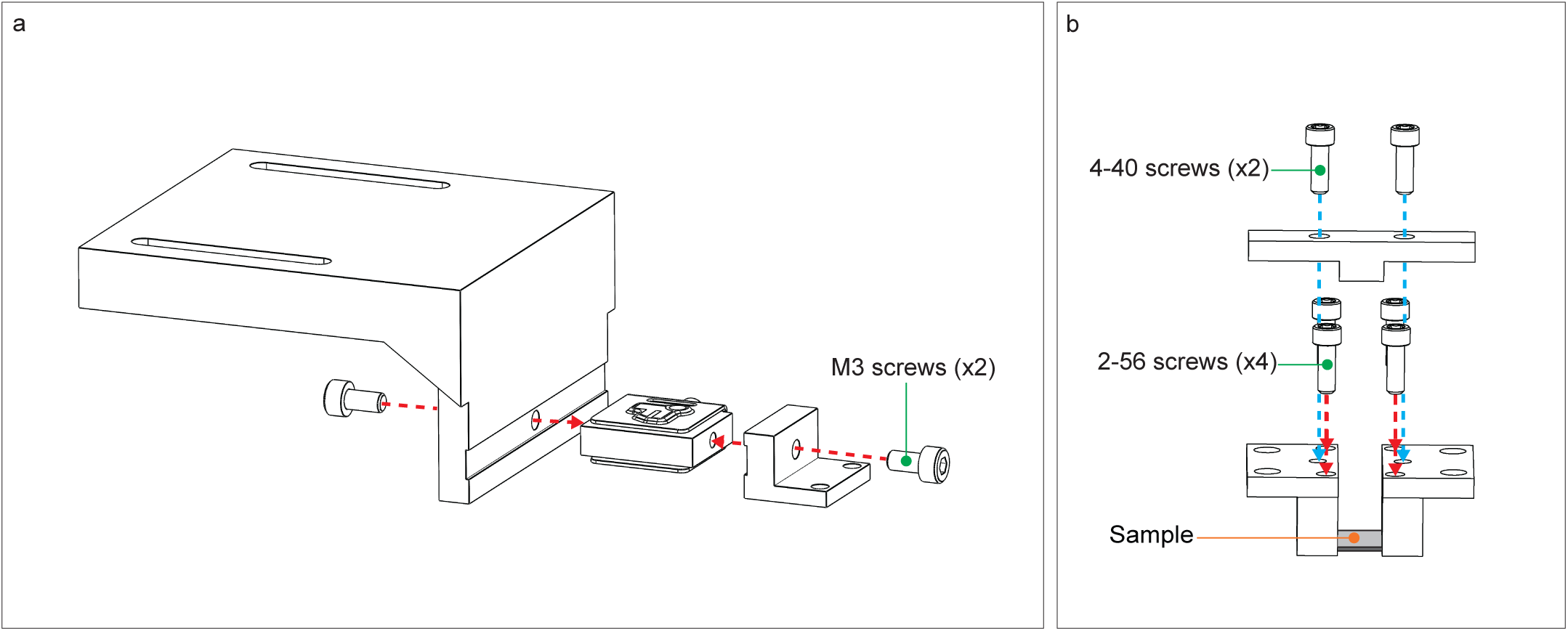
(a) Force transducer assembly. (b) Jig assembly. Arrows indicate location and direction of the screws for assembly. The location of the mounted sample being is indicated for reference.

### 5.3 Motor arms and jig

Parts required:

(*×*1) Motor arm

(*×*1) Motor arm with force transducer (Fig. 4a)

(*×*1) Jig assembly

(*×*4) 6-32 x 5/16” socket head cap screw

(*×*4) 4-40 socket head cap screw

Attach both motor arms (one with the force transducer attached from Fig. 4a and one without) to the top of each motor using 6-32 socket head cap screws through the arm slot and into the inner pair of screw holes on the motor closest to the imaging window (Fig. 5, red arrows). Position the motor arms so that the screws are at the end of the slot closest to the imaging window before tightening the screws. Suspend the assembled jig from Fig. 4b between the two motor arms and attach with 4-40 socket head cap screws (Fig. 5, blue arrows). Adjust spacing of motor arms as needed to fit the jig.

**Figure 5:**
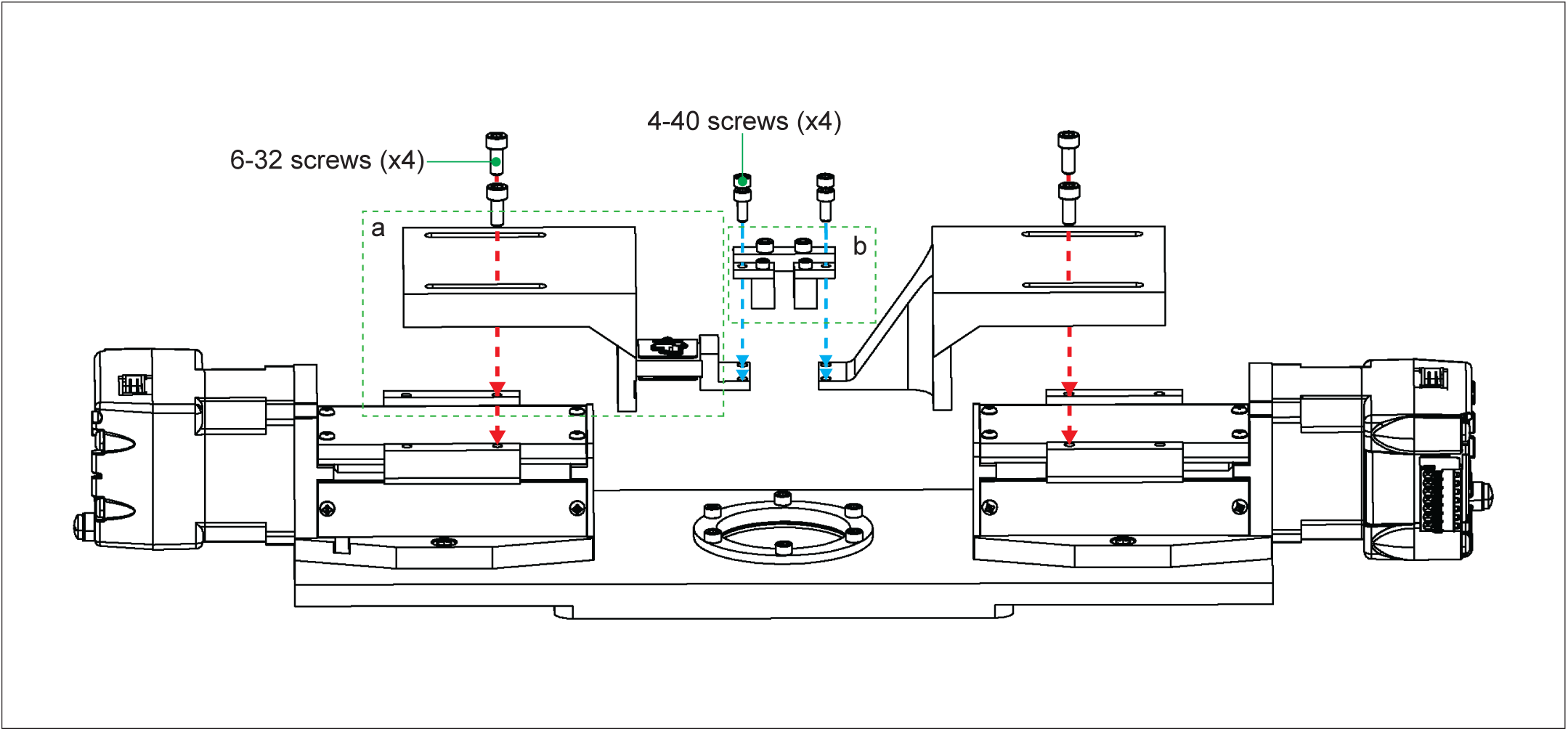
Steps to attach motor arms and jig to stepper motors to complete build assembly. Green dashed outlines refer to parts assembled in Fig. 4.

## 6. LabVIEW Control System Description

The control system of the stretcher was developed to provide a user-friendly interface to initiate stretching experiments, adjust experimental parameters, and conduct data collection. To achieve this, we employed the graphical programming environment LabVIEW (National Instruments) to integrate a graphical user interface (GUI) with communication to the motors and force transducer. LabVIEW was selected over other potential interface options such as a Jupyter Notebook due to the ease of creating a GUI and the existence of LabVIEW example code provided by the hardware manufacturers. Our final interface can be modified quickly and has ease of usability for users. Instead of running lines of code directly during operation, LabVIEW is displayed as buttons and tabs for an intuitive organized view.

The control system allows the user to send commands to the stretcher and receive data. Additionally, the control software ensures the adherence to mechanical limits of the stretcher, thereby safeguarding both the sample and the device from potential damage. Since stretcher is mainly for mechanical testing, we focused on developing the cyclic stretch control (Fig. 6), in which the user simply specified the stretch amplitude, cycle number, and frequency. Our device can reliably implement a frequency of up to 6.8 Hz at 10% strain, similar to the strain frequency found in heart or lung tissues [8]. During experiment, the software reports the displacements of both motors and the force transducer readout.

**Figure 6:**
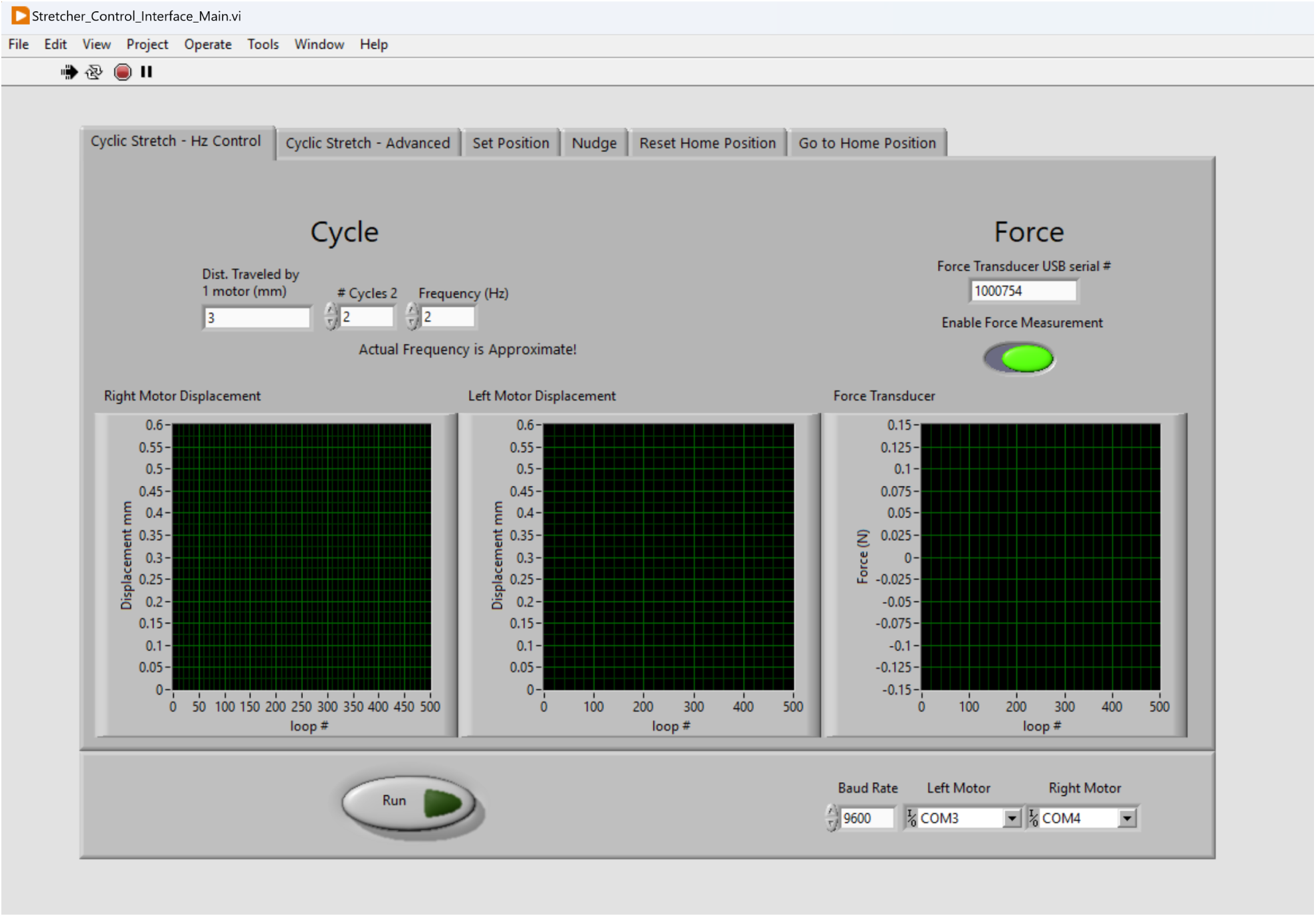
The Cyclic Stretch-Hz Control can be used for cyclic tensile testing as a simpler alternative to the Advanced tab, utilizing frequency (Hz) instead of velocity (mm/s).

As shown in Fig. 7, the block diagram of the control system is designed around an event structure that detects when the run button is pressed. This event triggers the execution of specific functions that are separated into tabs on the front panel. The control system employs a case structure to discern the selected tab and execute the corresponding functionality. During the execution of a linear stretching experiment, the control system automatically captures timestamped displacement and force data at a rate of 10 Hz. This data is then saved into a comma-separated values (.csv) output file. This approach ensures convenient data storage and can be directly inputted into MATLAB analysis for stress-strain curves and post-processing. The connection between the control system, motors, and force transducer is established using serial communication. A RS-485 to USB adapter bridges the connection between the motor and the computer. The control system is configured to match the default communication rate of the motor, which is set to 9,600 bits per second. The user can modify this communication rate using serial commands, and the control system can be adjusted accordingly using the GUI.

**Figure 7:**
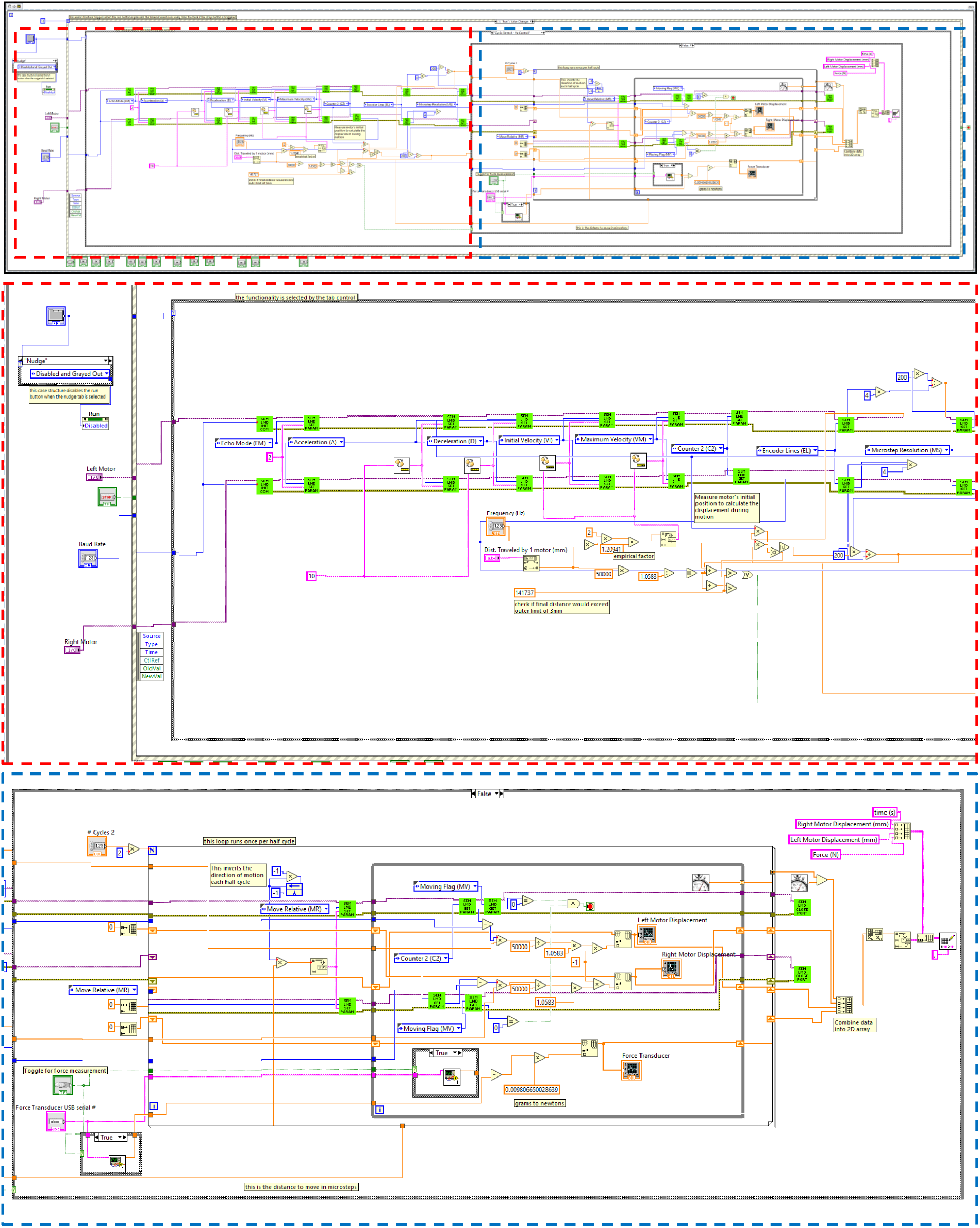
The block window breakdown view of the Cyclic-Hz Control tab.

We referenced the LabVIEW libraries provided by FUTEK (force transducer manufacturer) and Novanta IMS (stepper motor manufacturer) to write our code. We included the .net assembly FUTEK_USB_DLL.dll in the “get load cell measurement” sub VI which is necessary to communicate with the USB digital amplifier. We also utilized code from FUTEK’s LabVIEW example 13.0 for taking measurements from the force transducer. Similarly, sub VIs from Novanta IMS’s “LabVIEW Examples For LMD” are incorporated, allowing access to fundamental motor functions such as position measurement, movement control, and serial communication initiation and termination.

While the default waveform of cyclic stretch is seesaw, we also developed a Cyclic Stretch - Advanced function (Fig. 8), in which the users can define values of the motor acceleration, deceleration, initial velocity, and velocity upper bound, effectively changing the cyclic stretch waveform. The block window breakdown view of the Cyclic Stretch - Advanced function is summarized in Fig. 9. Both the Cyclic Stretch - Hz Control and Cyclic Stretch - Advanced tabs have the option to disable the force measurement by unchecking the Enable Force Measurement button.

**Figure 8:**
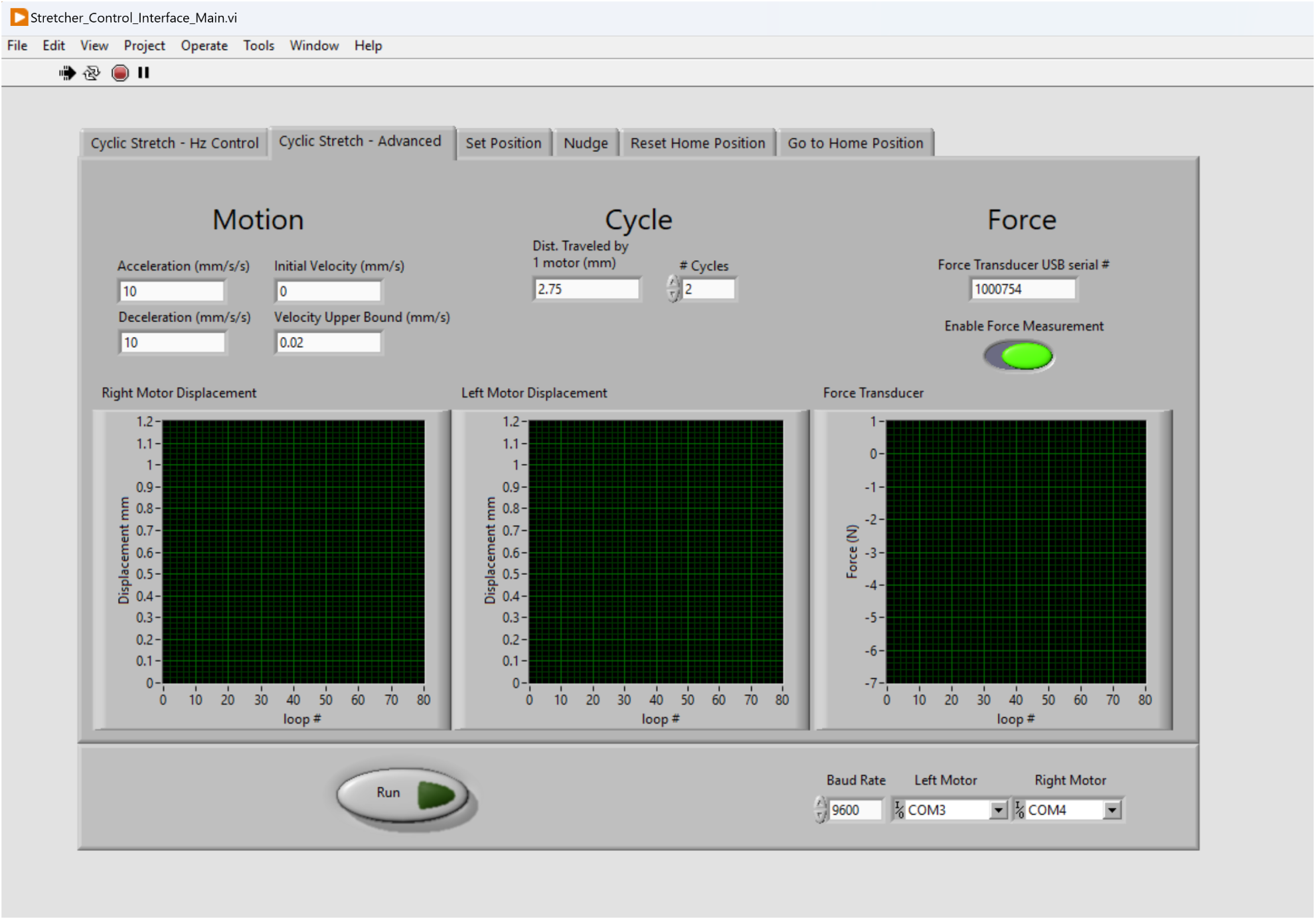
The Cyclic-Advanced tab can be used to conduct cyclic tensile testing with user-defined settings.

**Figure 9:**
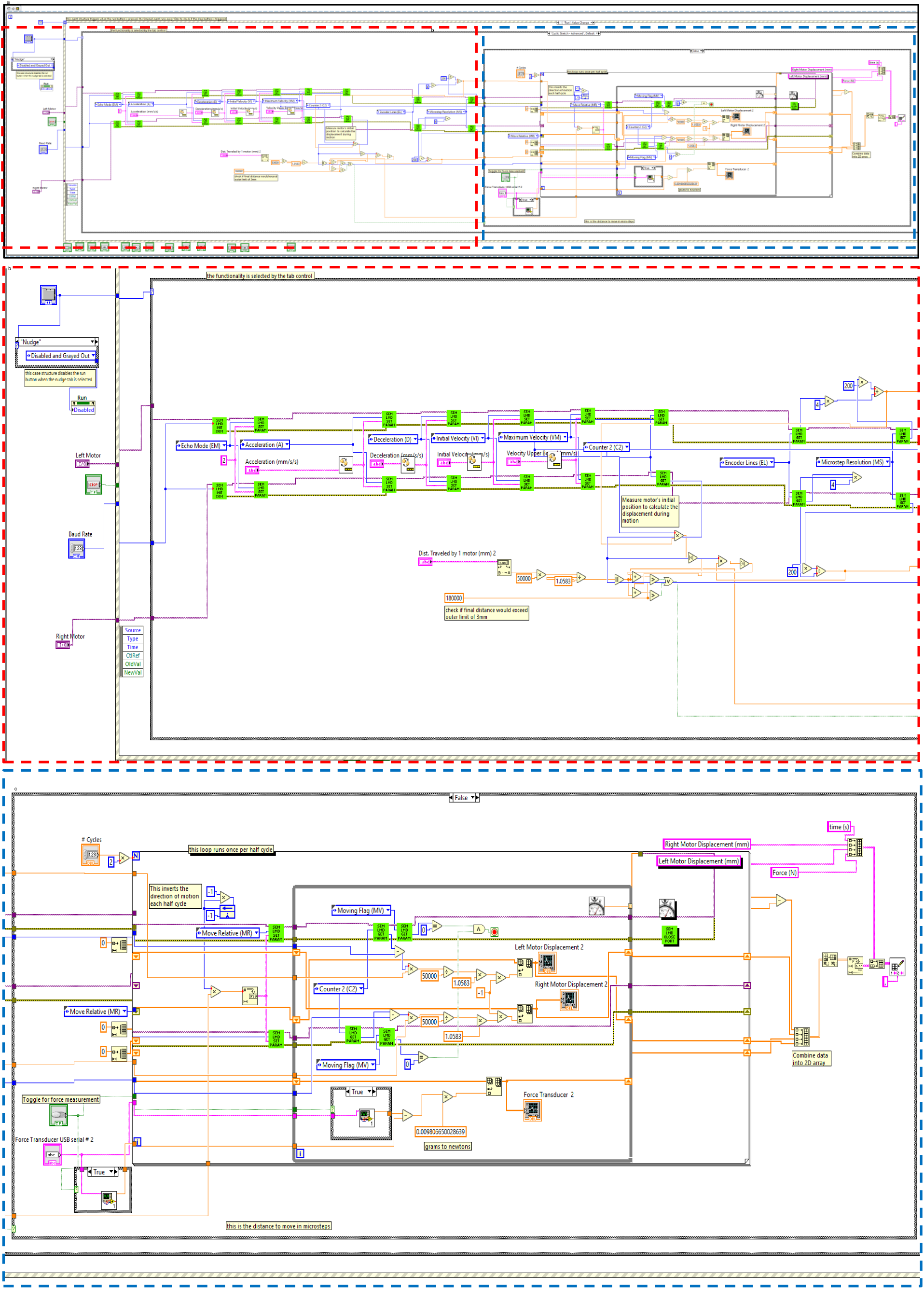
The block window breakdown view of the Cyclic-Advanced tab. Red dashed outline indicates the left portion of block code and the blue dashed outline indicates the right portion. The top row is both parts connected.

Since starting each experiment requires dismounting the old sample and mounting the new sample, it is essential to be able to automatically send the motors to a pre-set home position (Fig. 10). To do this, the users first use the Nudge tab (Fig. 11) to move the motors by 0.1 mm, 0.5 mm, or 1 mm incrementally inwards or outwards to the desired position, and subsequently use the Set Position tab to define the current position as the home position. The Nudge function is also useful for pre-stretching the sample to ensure the sample’s flatness.

**Figure 10:**
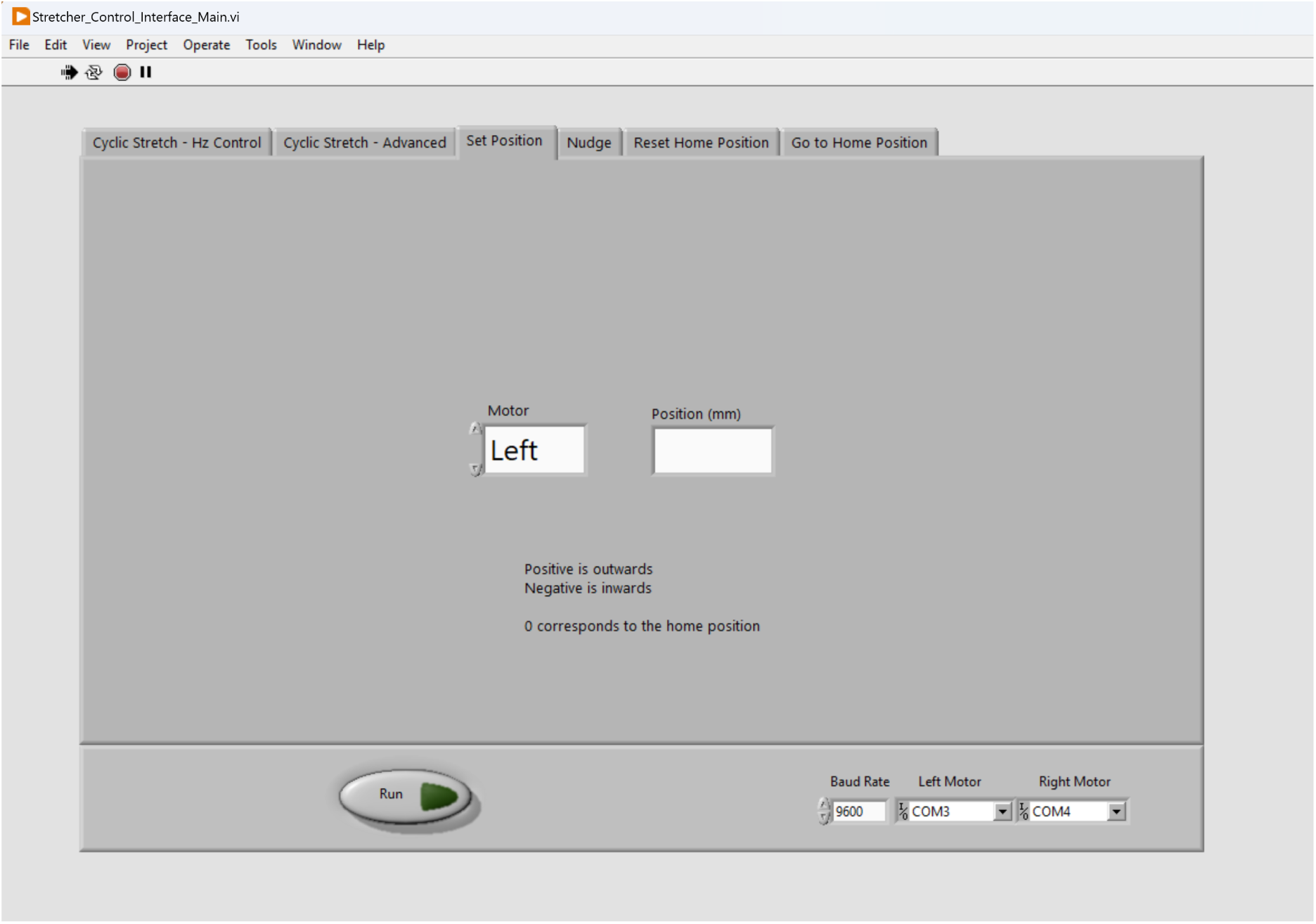
The “Set Position” tab can be used to set the current motor positions as the motors’ home position.

**Figure 11:**
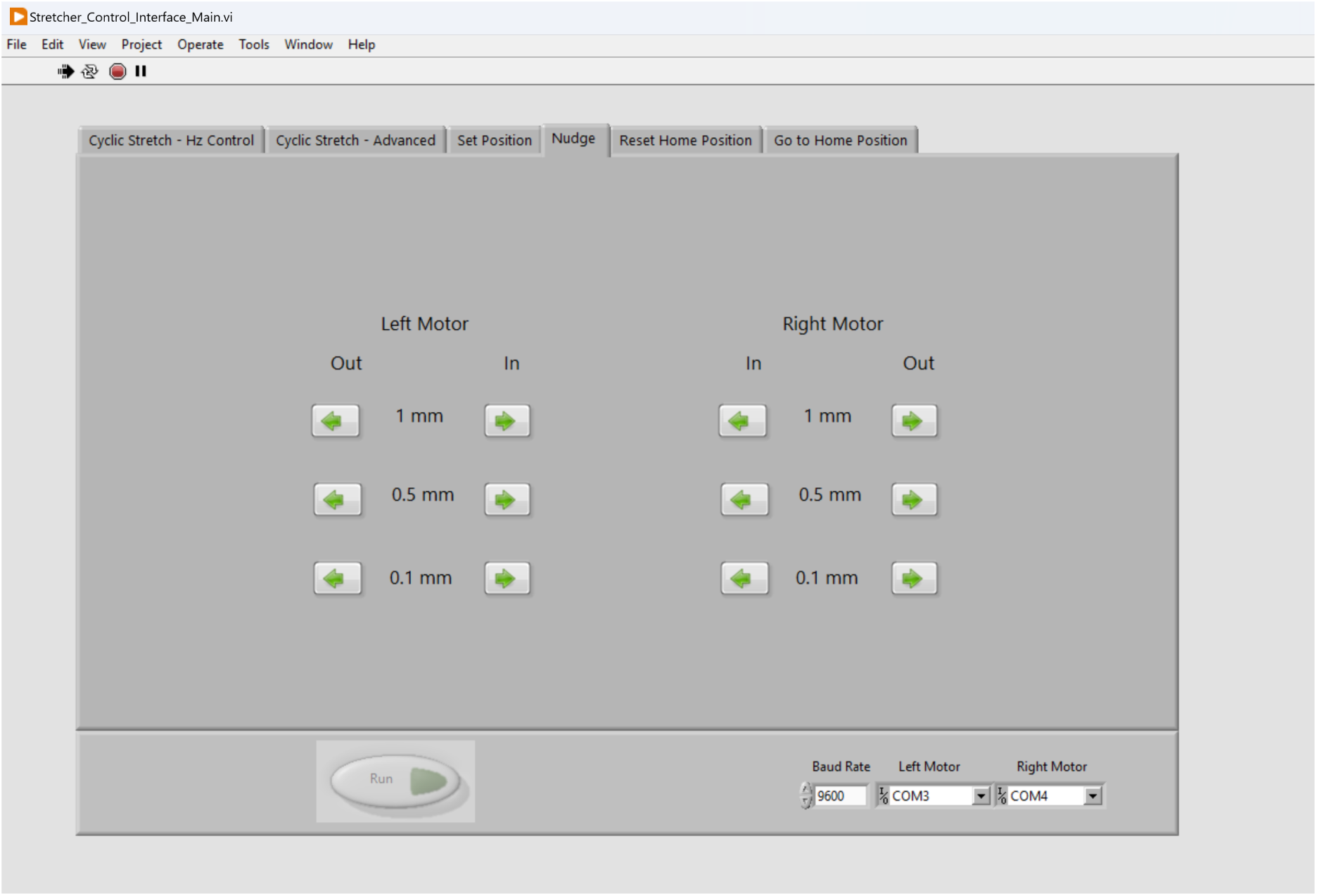
The “Nudge” tab can be used to move the motors by 0.1 mm, 0.5 mm, or 1 mm towards or away from the imaging window. This is useful for making sure suspended samples are tight when clamped into the jigs, allowing adjustment as needed.

We developed two methods to send the motors to the home position. The first method is to use the Reset Home Position tab (Fig. 12), which send both motors inwards to the limit switches (i.e., the most inner position), then outwards to the home position. When using this function, it is pivotal to ensure that no samples or jigs are attached to the arms before running. The second method to send the motors to the home position is using Go to Home Position, where both motors directly move to the home position. The Reset Home Position tab is useful if the users need to fully reset the motors.

**Figure 12:**
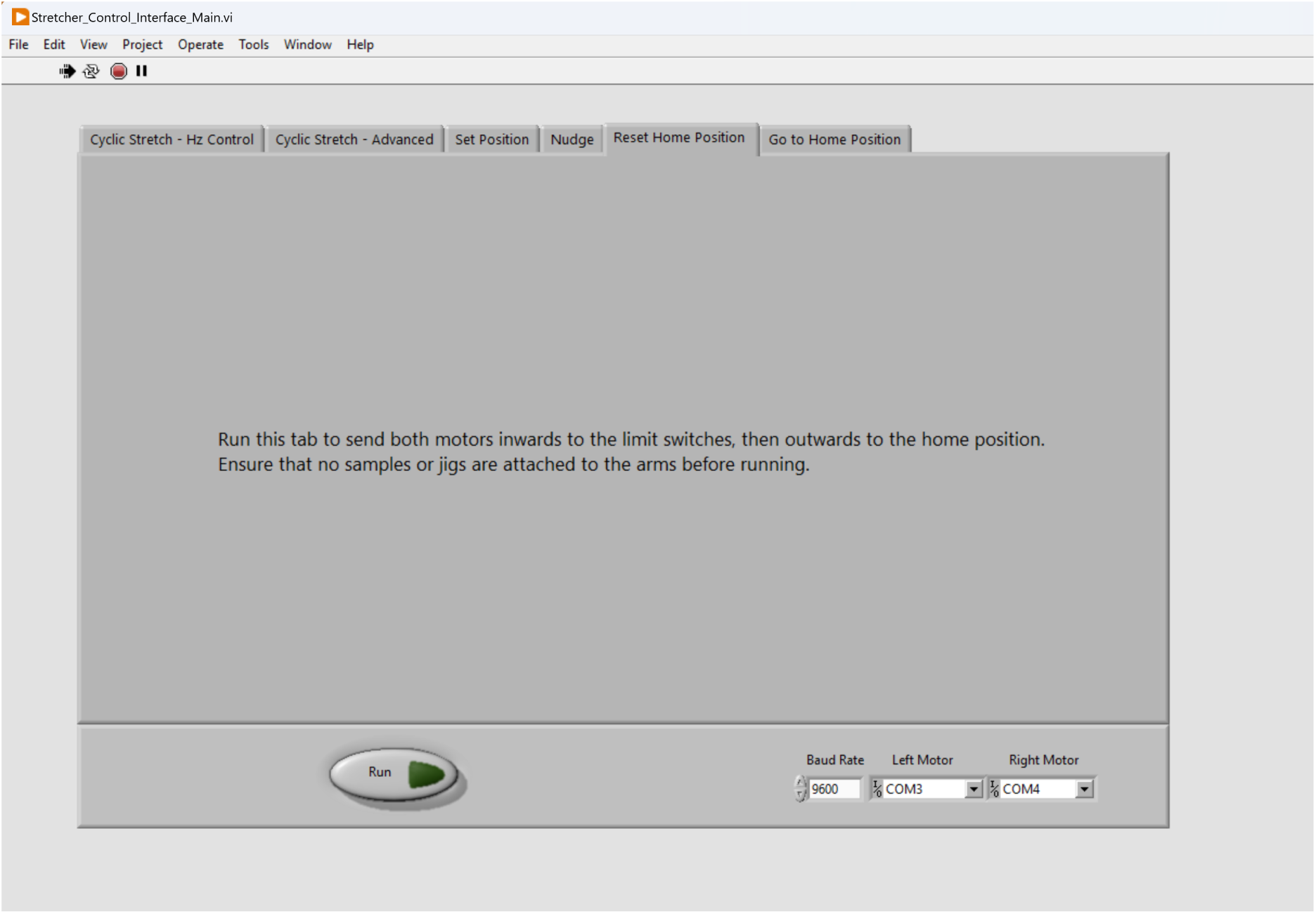
The “Reset Home Position” tab can be used to recalibrate the motors’ home position.

**Figure 13:**
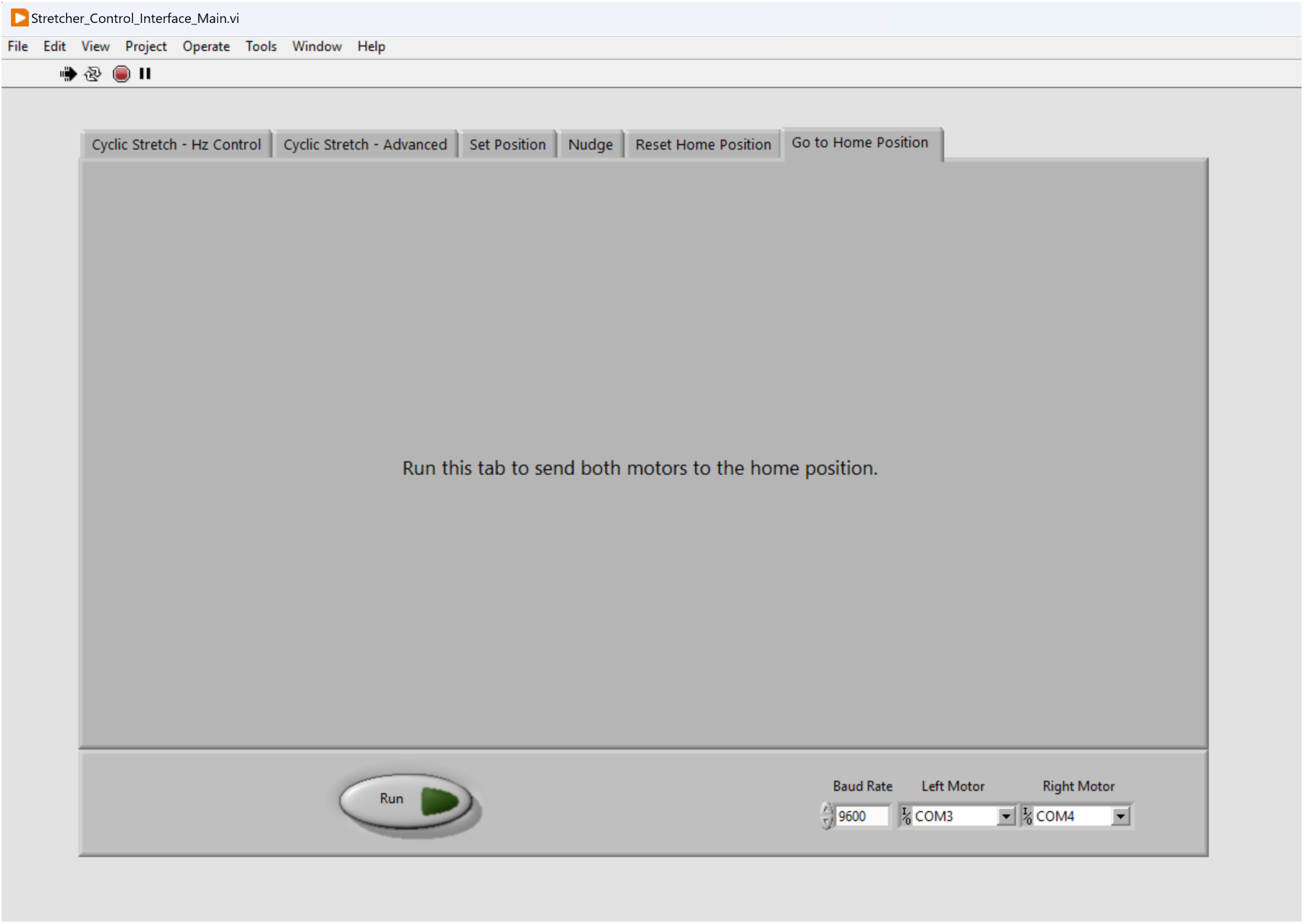
The “Go To Home Position” tab can be used to send the motors to the home position.

## 7. Operation instructions

In this section, we provide operation instructions using two common examples: particle-polymer membrane (silica particles embedded in polydimethylsiloxane, PDMS) and an epithelial cell monolayer cultured on a PDMS membrane. Specifically, we provide a step-by-step instruction of the sample preparation and mounting, as well as the software operation.

### 7.1 Sample Preparation and Mounting

We describe the sample preparation for both polymeric and biological samples. For the polymeric sample, we use PDMS/silica composites as an example as it is widely used due to its tuneable stiffness and bio-compatibility. For the biological sample, we used an epithelial cell monolayer as an example as it is commonly used in biomechanics and mechanobiology studies. It should be noted that while the cell substrate must form a free standing membrane spanning the length between the jig legs to enable sample stretching, the cells themselves do not need to form a fully confluent monolayer. For *ex vivo* samples where tissues are directly obtained from a host, such as a mouse model, no fabrication is required. The tissue can simply be trimmed to the desired size and directly clamped to the jig using the mechanical clamps previously described. If the sample exhibits slacking after mounting, we recommend correcting for the slack using microscopy. Slacking will be illustrated by out of focus edges due to differences in sample height resulting from sample sagging. If this is observed, we advise to correct for this by slowly stretching the sample while performing live imaging until tautness is achieved once the full field of view is in focus.

#### 7.1.1 Polymeric Sample

1. Coat your desired oven-safe sample mold with a thin layer of water-soluble polyvinyl alcohol (PVA). This coating allows sample to be easily removed from the mold in subsequent steps. As an example, we use an 8 mm x 15 mm x 2 mm mold size. Recommended sample thickness and length is 1 mm and minimum 15 mm, respectively.
2. Place PVA-coated mold in an oven or on hotplate at 100 *^◦^*C for PVA to dry. The PVA is dry when no visible liquid remains in the mold.
3. In a weigh boat, measure out desired mixing ratio of Sylgard 184 base and curing agent and mix evenly for 2-3 minutes. If using particles, weigh out particle amount and combine, mixing for another 2-3 minutes or until fully incorporated. Degas weigh boat for 7 minutes using a desiccator, ensuring no air pockets remain. Here, we use Sylgard 184 as an example, but other formulations or polymers may be utilized.
4. Pour mixture into mold using scalpel edge to scrape the top and make samples level. Ensure there are no air pockets from pouring. Use a manual hand-held air pump to pop any remaining bubbles if present. Cure sample at 75 *^◦^*C for 24 hours.
5. Run mold under water to dissolve PVA coating then use scalpel to carefully cut out samples from mold.
6. For staining, submerge samples in 1:250 BODIPY to deionized water solution for 15 hours. Cut to size for imaging and mounting. 5 x 15 x 1 mm strips are recommended. Here, we use BODIPY as an example stain for lipophilic compounds.
7. To mount, invert the jig with the legs assembled to the T-bar so that the bottom of the jig legs face up. Lay the sample strip perpendicular to the legs, bridging across the two sides as straight as possible (Fig. 2b).
8. Slide the jig clamps onto the jig legs (Fig. 2b), pressing into the sides of the sample and attach the clamps to the legs with 2-56 screws (Fig. 4b), avoiding relaxation of the suspended sample. Mount prepared jig to the motor arms as detailed in building instructions Section 5.3.

#### 7.1.2 Biological Sample

1. Spin coat a clean glass coverslip with PVA at 2000 RPM for 2 minutes and cure for 5 minutes at 150 *^◦^*C.
2. Spin coat desired mixing ratio of Sylgard 184 at 2000 RPM for 2 minutes on cured PVA coating from Step 1. Spin speed and duration should be optimized based on desired sample thickness.
3. Apply a thin layer of uncured Sylgard 184 mixture to the bottom of the assembled jig (Fig. 4b) and gently place the jig on the PDMS-PVA coated coverslip such that the bottom of the jig legs are firmly adhered to the substrate. Cure the jig-coverslip composite for 35 minutes at 150 *^◦^*C.
4. Place jig in a glass petri dish and autoclave before culturing cells directly on the PDMS. A coating of fibronectin before seeding may be necessary to facilitate cell attachment to PDMS.
5. Culture cells as desired. When ready to stretch, remove the jig-coverslip composite from the petri dish and gently cut excess PDMS not enclosed between the two jig legs using a sterilized scalpel. Mount freestanding membrane suspended between jig legs to stretcher as described in building instructions Section 5.3. Sample should be firmly suspended between jig legs such that jig clamps are not required.

### 7.2 Software Operation

1. Install LabVIEW on a computer.
2. Move the provided FUTEK_USB_DLL.dll into the file directory containing LabVIEW.exe.
3. Connect the stretcher control box to the stretcher using the two ribbon cables, two barrel plugs, and force transducer connector contained within the braided cable sleeve.
4. Connect the stretcher control box to the computer via the included USB cable. The stretcher control box should only be connected to USB 3.0 ports (blue) to prevent damage to the force transducer.
5. Connect the stretcher control box’s two AC plugs to a power source.
6. On the connected computer, run Stretcher_Control_Interface_Main.vi to open the virtual instrument (VI).
7. Run the VI by pressing Ctrl+R or Cmd+R on the keyboard.
8. Use the “Left Motor” and “Right Motor” dropdown buttons found on the bottom panel of the VI to select the corresponding communication ports.
9. The functionality of the user interface is divided into tabs, which can each be executed using the “Run” button found in the bottom panel of the VI:

(a) Reset Home Position: Run to recalibrate the home position. Ensure that no samples or jigs are attached to the arms before running.
(b) Go to Home Position: Run to send both motors to the home position.
(c) Nudge: Use this tab to move each motor in incremental distances of 1 mm, 0.5 mm, or 0.1 mm.
(d) Set Position: Run to send a motor to a position relative to the home position.
(e) Cyclic Stretch - Hz Control: Run to conduct cyclic stretching. Ensure motors are either set to home position or starting at zero displacement using the Set Position tab. Motor speed is calculated by the selected frequency and stretching amplitude. If using a force transducer, input the serial number of the USB digital amplifier into the box provided.
(f) Cyclic Stretch - Advanced: Run to conduct cyclic stretching. Ensure motors are either set to home position or starting at zero displacement using the Set Position tab. Motor speed is determined by selected motion parameters. If using a force transducer, input the serial number of the USB digital amplifier into the box provided.
10. After running a cyclic stretching experiment, select a location to save the output data file.
11. Stop the VI using the red octagon button (Abort Execution) found in the top panel of the VI.
12. Disconnect the stretcher control box AC plugs.
13. Disconnect the stretcher control box USB.

Tips: Always connect and disconnect the stretcher’s cables in the order described above to prevent damage to the electronics. Avoid plugging or unplugging cables while the motors are moving. To avoid unexpected behavior, do not interact with the user interface while the motors are moving.

## 8. Validation and characterization

The central novelty of our stretchoscope is the integration of the motion control, force measurement, and microscopy imaging during membrane stretching. Therefore, it is essential to validate and characterize each of the three functions. For the motor displacement characterization, we determined its repeatability to be less than 3.5 *µ*m, corresponding to a tensile strain *∼* 0.03%. For force measurements, we also quantified the force readout repeatability as *∼* 60 mN. We further confirmed the force measurement correctness by directly comparing the our measured Young’s moduli with that measured by Instron, a leading mechanical testing machine. Lastly, we provided two imaging examples to illustrate practical workflows of combining the stretchoscope, fluorescent microscopy, and image analysis. In the first demonstration, we showed the plane strain field in an uniaxially stretched silica/PDMS composite. In the second demonstration, we showed the cellular deformation in a uniaxially stretched epithelial cell layer.

### 8.1 Motor Displacement Consistency

The motor displacement consistency during cyclic stretching was investigated by tracking the edge of a mounted jig leg and evaluating its displacement amplitude (Fig. 14a). To do this, we mounted the stretchoscope on a standard inverted light microscope (10*×* NA=0.25 objective) and video recorded (30 frames per second) 25 cycles of stretching. While each image pixel corresponds to *∼*0.26 *µ*m, the actual imaging resolution is *∼*1.5 *µ*m, which is mainly determined by the objective NA=0.25. Example images of the jig at representative positions are shown in Fig. 14a. The captured images were then analyzed to identify the highest contrast line as the edge of the jig. The motion consistency was defined as the repeatability of the cyclic stretch amplitude between individual cycles, in which the amplitude was defined as the vertical distance between the peak and trough of a displacement cycle. We conducted three trials for each motor and plotted the displacement profile (Fig. 14b), the amplitude as a function of cycle number (Fig. 14c), and the histogram of the amplitudes (Fig. 14d). The displacement profile (Fig. 14b) confirms the seesaw waveform of the cyclic stretch as defined in the control software. The displacement amplitude analysis also confirms little to no drift of the amplitude over time. Lastly, we observed a unimodal probability distribution of the amplitude for both motors. By calculating the standard deviation, we found that the left motor’s repeatability (one *σ*) was within 3.5 *µ*m and the right motor was within 2 *µ*m. We note that our measured motor motion consistency may over-estimate the true value since our microscopy system (10*×* objective NA=0.25) can only achieve a micron-scale resolution. Nevertheless, the 3.5 *µ*m displacement repeatability corresponds to a tensile strain repeatability *∼* 0.03%, which well exceeds the requirement of typical mechanical tests.

**Figure 14:**
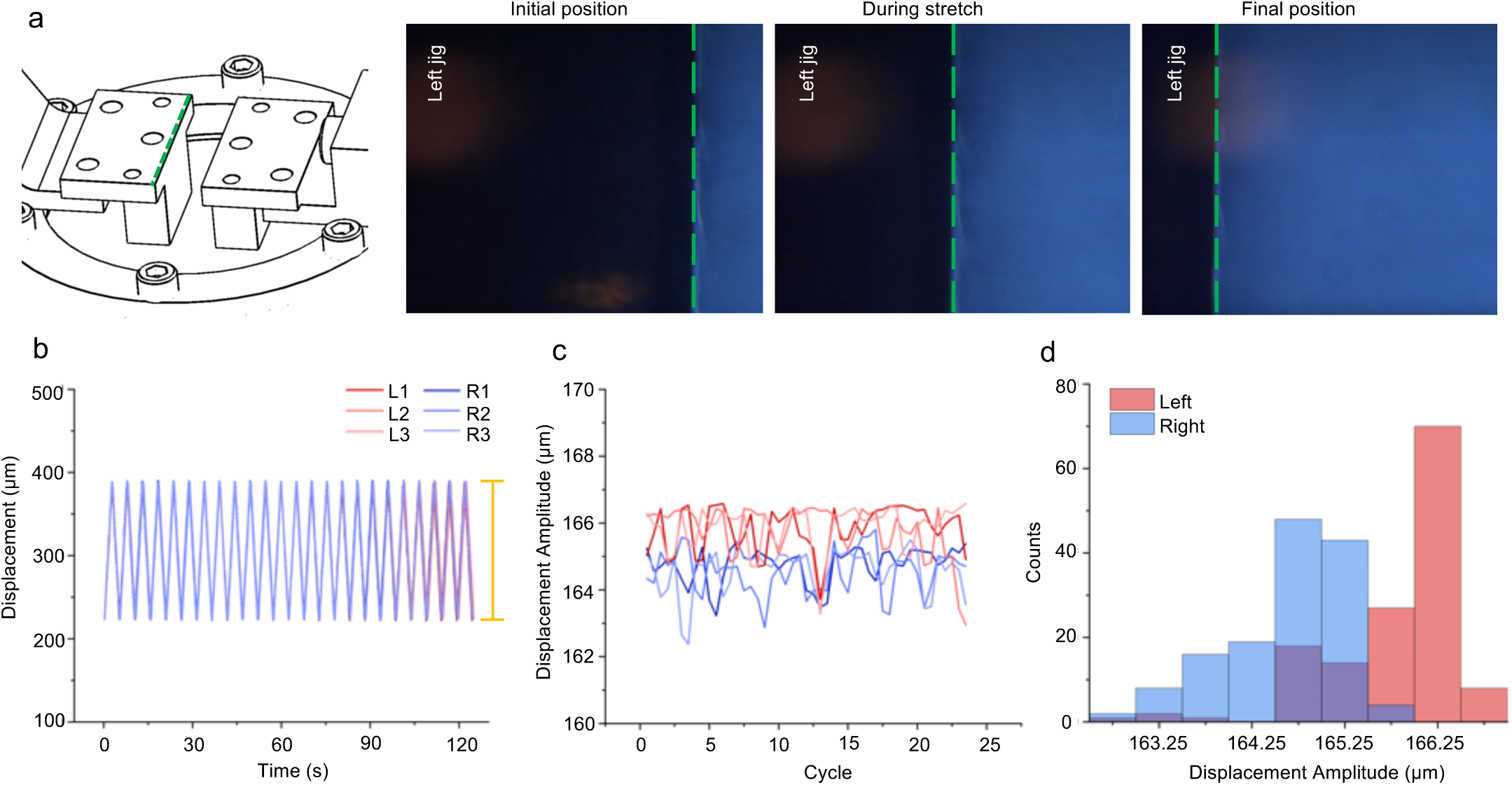
Motor motion repeatability analysis. (a) Schematic of jig being stretched. Green dashed line indicates edge being imaged for motor motion validation. From left to right, snapshots of left jig leg moving through initial, middle, and end positions in video being analyzed. (b) Displacement of jig edge vs. time for left (L) and right (R) motors across three trials of 15 cycles for each side. Displacement amplitude from maximum to minimum of cycles indicated by yellow bar. (c) Displacement amplitude values vs. cycle for left and right motors. Legend same as Fig. 14b. (d) Probability distribution of displacement amplitudes for left and right motor.

### 8.2 Force Transducer Consistency

The force transducer was calibrated by the manufacturer using a 5-point calibration test in both tension and compression (Fig. 16). If a manual calibration is desired, a scale method calibration can be done using the force transducer’s SENSIT software and the protocol outlined in the SENSIT Software Manual. The force transducer consistency was evaluated by stretching an elastomeric strip (thickness *∼* 0.9 mm, width *∼* 4.5 mm) with uniform mechanical properties and evaluating the force amplitude. We mounted a 7.45 mm segment of elastomeric strip onto the stretchoscope. We ran 50 cycles at 20% strain and analyzed the force profile (Fig. 15a), amplitude as a function of cycle number (Fig. 15b), and the amplitude histogram (Fig. 15c). As described in the LabVIEW Control System Description section, the force readout was automatically recorded and saved as a .csv file. Similar to the motor displacement analysis, the force amplitude was defined as the vertical distance between the peak and trough of a force cycle. This experiment also allows us to assess the displacement consistency with the motors under load. To do this, we simultaneously monitored the motor displacement by reading the motor’s encoder. Using such data, we also analyzed the displacement profile (Fig. 15d), amplitude as a function cycle number (Fig. 15e), and the amplitude histogram (Fig. 15f).

**Figure 15:**
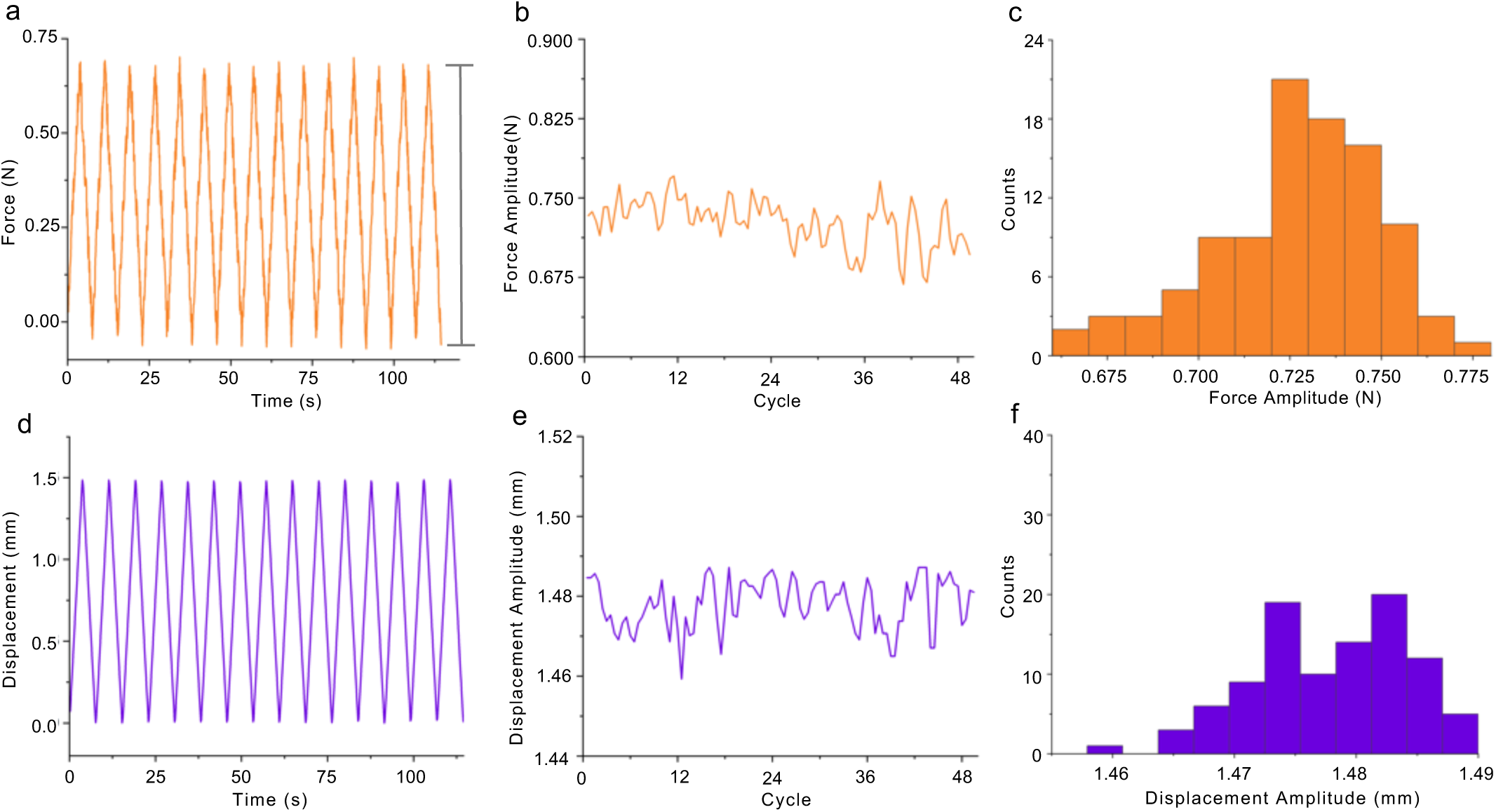
Force measurement repeatability analysis. (a) Force vs. time plot of elastomer over the first 15 cycles (not all cycles shown to allow for amplitude visualization). Force amplitude from maximum to minimum of a cycle indicated by gray bar. (b) Force amplitude over 50 cycles. (c) Probability distribution of force amplitude for all 50 cycles. (d) Displacement vs. time plot of elastomer over the first 15 cycles (not all cycles shown to allow for amplitude visualization). (e) Displacement amplitude over 50 cycles. (f) Probability distribution of displacement amplitude for all 50 cycles.

**Figure 16:**
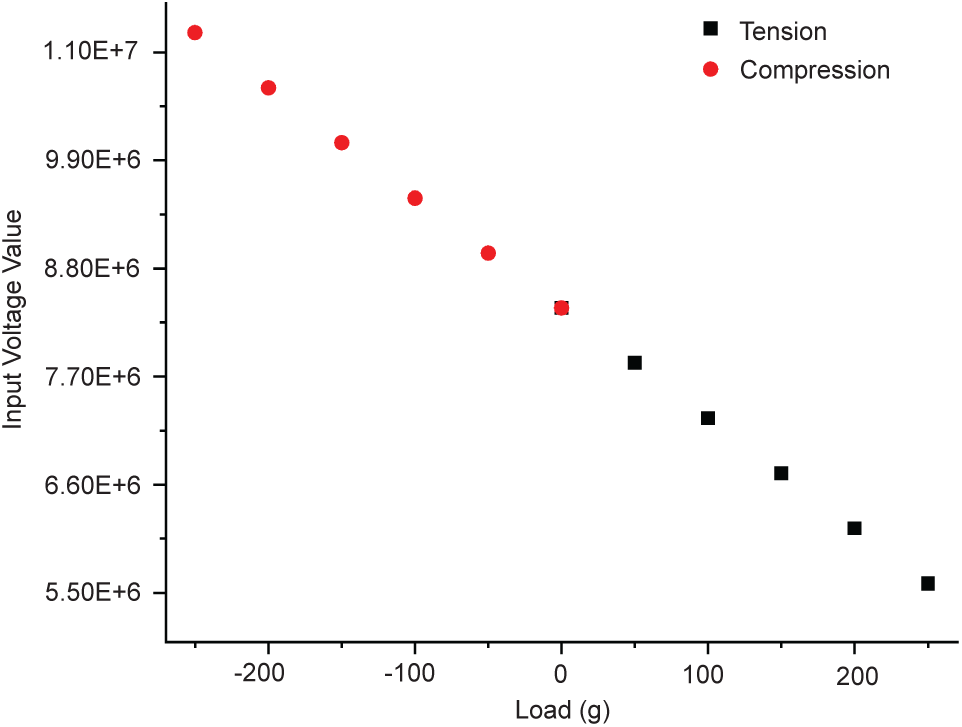
Calibration curve for force transducer from manufacturer (FUTEK). Five different loading weights were used in each loading direction (tension and compression). The zero point was also measured separately for each loading direction.

As shown in Figs. 15 a and d, both the force and displacement profiles exhibit seesaw waveforms, as defined in the control software. Furthermore, we did not observe significant drifts in either the force amplitude or displacement amplitude over time. By calculating the histogram standard deviation, we found that the force consistency was within 0.06 N (*<*8% of mean), whereas the displacement consistency was within 30 *µ*m (*<* 2% of mean). These results suggest that both the force and displacement measurements are reproducible between testing cycles.

### 8.3 Linear Stretching Validation

To assess whether the stretchoscope can accurately determine the mechanical property of the sample, we conducted a parallel modulus measurements using our device and the commercial Instron 5943 with a 50 N load cell attached. The tested samples are PDMS (Sylgard 184) strips with two different ratios of PDMS curing agent to base, 1:40 and 1:25. For each condition, three samples were run on each device. We found that the stress-strain curves produced by the stretchoscope agrees with that from Instron 5943 for both the 1:40 (Fig. 17a) and 1:20 (Fig. 17b). The linear modulus of each sample was then determined as the mean slope of the 0% to 5% strain region. Notably, while the stretchoscope stress-strain curves appeared noisier than the Instron data, such a stress fluctuation did not significantly impact the linear modulus measurements, which was determined based on linear regression. For the 1:40 samples, the stretchoscope modulus was 0.0375 MPa and the Instron modulus was 0.0251 MPa (Fig. 19c). For the 1:25 samples, the stretchoscope modulus was 0.2429 MPa and the Instron modulus was 0.2102 MPa (Fig. 19c). The apparent modulus difference between the 1:40 and 1:20 samples for both the stretchoscope and Instron measurements indicate that our device can discern mechanical property differences between the two tested conditions.

**Figure 17:**
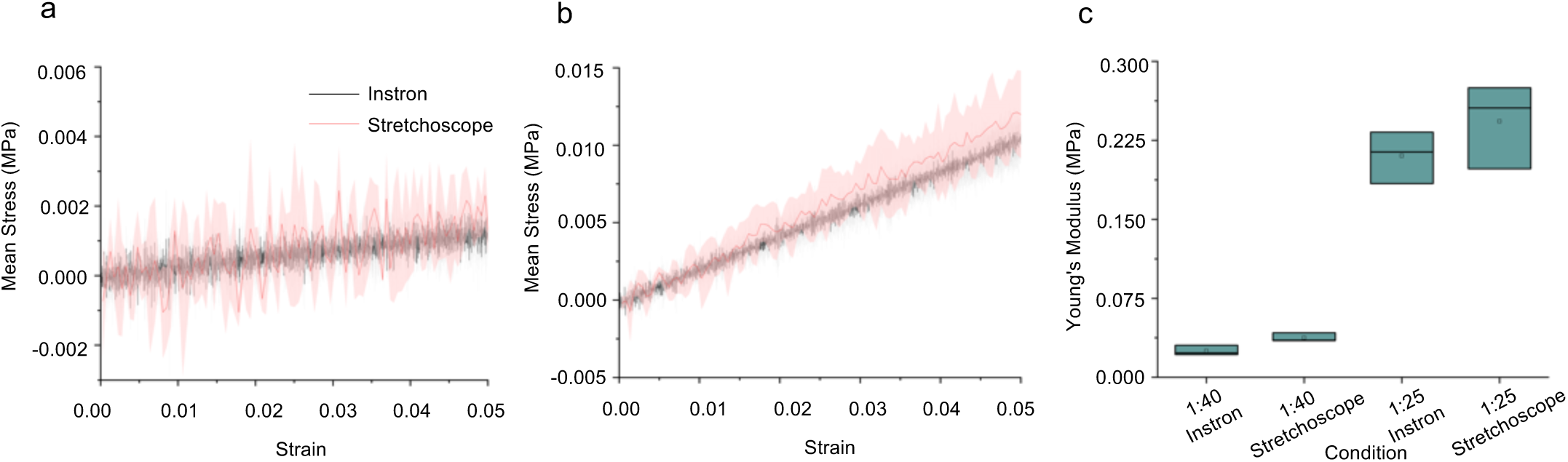
Validation of elastic modulus measurements. (a) Stress strain curves of 1:40 neat PDMS samples measured using Instron (black) and the stretchoscope (red). Solid line and shaded area represent the mean and standard deviation, respectively. (b) 1:25 stress strain curves, solid line and shaded area are mean and standard deviation, respectively. (c) Box plot showing the linear elastic moduli of 1:40 and 1:20 PDMS samples characterized using Instron and the stretchoscope.

### 8.4 Strain Field Uniformity Analysis

The strain field uniformity was evaluated using Particle Image Velocimetry (PIV) [23]. Samples were created using the procedure in Section 7.1.1 on polymeric sample preparation. The ratio of base to curing agent for the PDMS was 1:40 and the PDMS was combined with 150% silica dioxide particles by weight. After curing, the sample was then stained with an oil-based fluorescent dye (BODIPY). The stained sample was then mounted on the stretchoscope and imaged using a confocal microscope (RCM1 on Nikon Ti-E) with a laser excitation at 488 nm. The sample was imaged before (Fig. 17a) and after (Fig. 17b) the uniaxial stretch. Since BODIPY selectively stains the PDMS, the PDMS and silica particles appear bright and dark in the confocal image, respectively.

The confocal images were then processed using the PIVlab add-in for MATLAB, creating the displacement field (Fig. 17c). The analysis used two passes. The parameters for the first pass were an interrogation window of 150 and step size of 35. The parameters for the second pass were an interrogation window of 20 and step size of 10. This displacement field was used to calculate the fields of *x* displacement (Fig. 18d), *y*-displacement (Fig. 18e), *XX* strain (Fig. 18f), and *yy* strain (Fig. 18g). Here, the *xy* plane is the plane of the sample face, in which the *x*-direction is the stretch direction, and the *y*-direction is its normal axis. As shown by Figs. 17d and 17e, we found that the displacement fields are relatively uniform, as expected from a tensile stretch. As anticipated, we observed positive values for the *xx* strain component (mean *∼* 0.075), and negative values for the *yy* strain component (mean *∼* -0.03). These results demonstrate that the stretchoscope can be integrated with confocal microscopy and image-based strain analysis to characterize microstructural responses in stretched materials.

**Figure 18:**
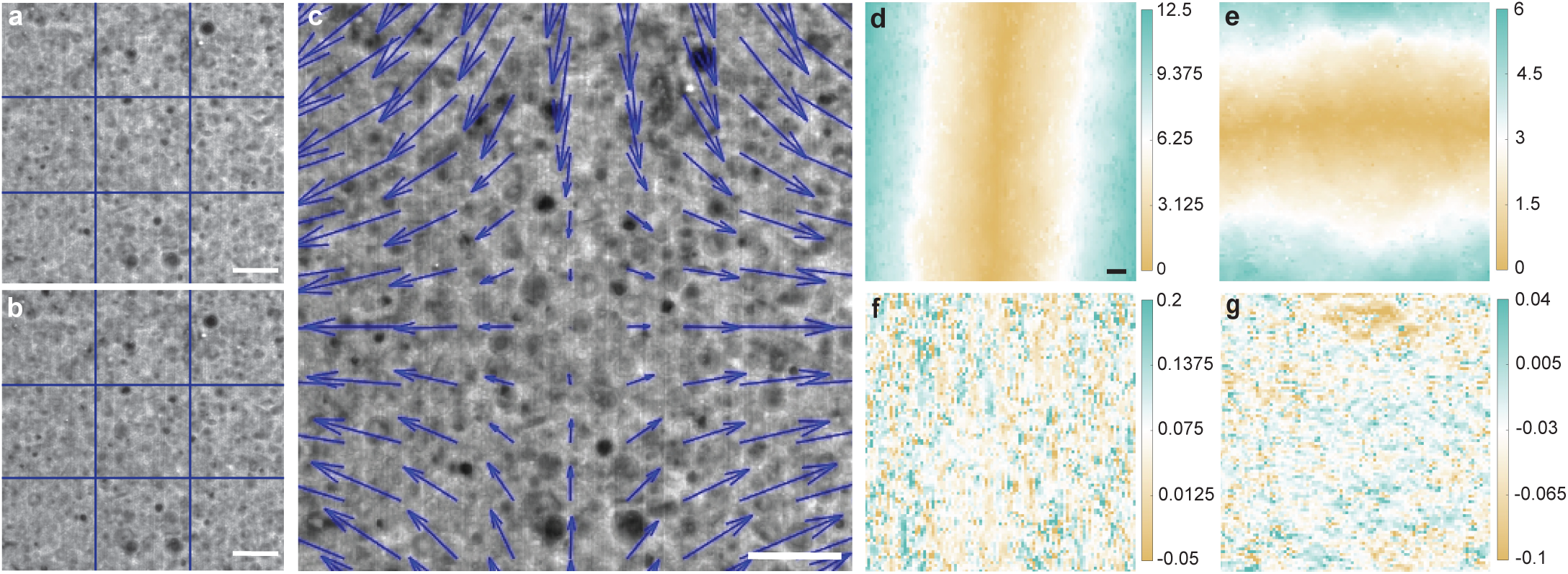
Strain field uniformity analysis. (a) 1:40 PDMS sample doped with silica particles before stretch. (b) Same reign of interest shown in (a) after stretch. (c) displacement field generated by PIVlab demonstrating the shift of particles with 20% strain. (d) Absolute displacement for *x*-direction. (e) Absolute displacement for *y*-direction. (f) Strain *xx* distribution (unitless). (g) Strain *yy* distribution (unitless). All scale bars are 20 *µ*m. Grid lines in (a) and (b) are added as a guide for eyes.

### 8.5 Imaging of stretched cell monolayers

Cell samples were prepared using Madin-Darby canine kidney (MDCK) cells following the procedure in Section 7.1.2 on biological sample preparation. The ratio of curing agent to base for the PDMS was 1:10. The jig was mounted on the stretchoscope and imaged using a confocal microscope (RCM1 on Nikon Ti-E) with a 20*×* NA=0.75 objective. The sample was imaged before and after a 50% tensile strain (Fig. 19). By comparing the unstretched and stretched images, we observed elongation of the cell shape due to the applied tensile strain.

**Figure 19:**
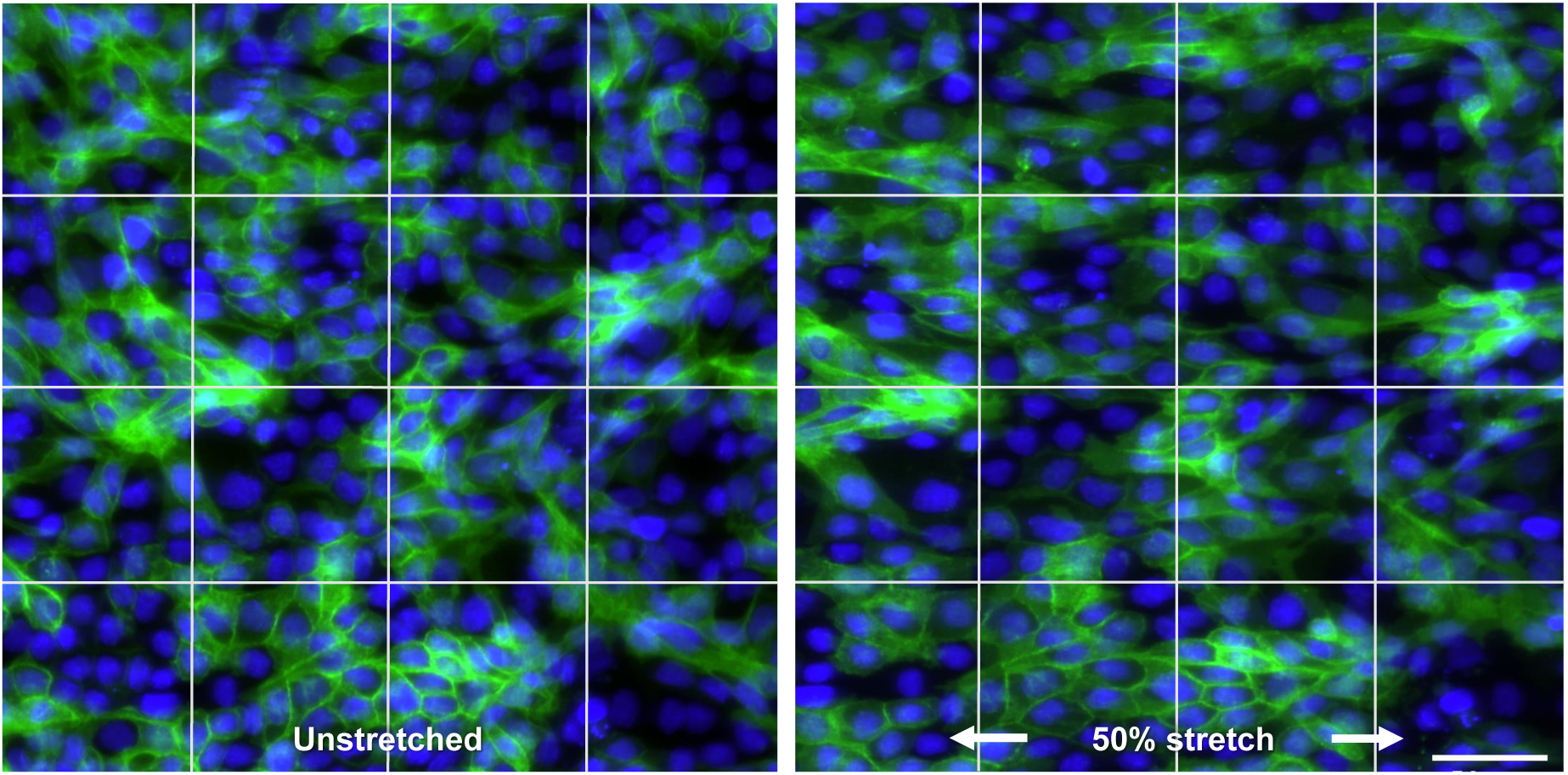
Comparison of unstretched (left) and 50% stretched (right) Madin-Darby Canine Kidney (MDCK) cells in the *x*-direction (horizontal). Green channel indicates plasma membrane labeled by green fluorescent protein (GFP), blue channel indicates nuclei labeled by Hoechst. Scale bar is 50 *µ*m. Grid lines are for displacement visualization purposes.

## 9. Conclusion

In this work, we report the development of a uniaxial membrane stretcher, stretchoscope, that allows simultaneous force measurement and inverted microscopy imaging. The force measurement of the stretchoscope provides a performance comparable to a commercial tensile testing device while also allowing for high resolution imaging. It has a high degree of motor displacement and force reading consistency and creates uniform strain fields. Some differences between our instrument and commercially available instruments, such as Instron, that may contribute to our slightly noisier data may be attributed to features such as material handling, sensitivity of measurement components, and software optimization. Using Instron, samples are mounted vertically in which the effects of sagging are negligible since the sample is oriented perpendicular to gravity, which is not feasible in our setup since our platform couples stretching with microscopy. In addition, many Instron instruments feature customized grips that can be optimized depending on sample material, shape, and size. Furthermore, Instron uses an AVE2 strain measurement device that can measure changes down to 0.5% of the reading. It is also possible that the Instron features technology to aid in minimizing external vibrations. To make our device more accessible financially, we have opted to utilize more economic devices that still provide accurate measurements. If future users hope to improve the measurement precision and accuracy, they can consider upgrading some components, such as the motors or force transducer, or add in an additional analysis step that can correct for external vibrations.

In addition to providing measurements comparable to commercially available instruments, our device is highly accessible and straightforward to assemble. The assembly of this device only requires standard hand tools and thus can be completed in any undergraduate-level mechanical laboratories. The LabVIEW GUI provides an intuitive and integrative way to control the stretcher and has a wide variety of options for control. Our published design files further enable the option to 3D print the device, making the instrument affordable and accessible. Overall, our stretchoscope serves as a valuable mechanical-structural experimental tool for both materials sciences and biomedical studies.

## CRediT author statement

***Shannon Li****: Methodology, Formal analysis, Investigation, Validation, Writing - original draft.* ***Alyssa Gee****: Methodology, Formal analysis, Investigation, Validation, Writing - original draft.* ***Nathan Cai****: Methodology, Software, Investigation, Writing - original draft.* ***Alexandra Bermudez****: Validation, Investigation, Supervision, Writing - review & editing.* ***Neil Y.C. Lin****: Conceptualization, Methodology, Investigation, Funding acquisition, Writing - review & editing*.

## Acknowledgements

The authors thank Lihua Jin, Chen Wei, and Boliang Wu for Instron training and generously allowing us to use the instrument. The authors also thank Jerry Chen for obtaining the MDCK cell stretching images. Funding: This work was supported by the UCLA SPORE in Prostate Cancer Grant [P0 CA092131] and National Science Foundation [CMMI-2029454].

